# BMAP: a comprehensive and reproducible biomedical data analysis platform

**DOI:** 10.1101/2024.07.15.603507

**Authors:** Yongyong Ren, Zhiwei Cheng, Leijie Li, Yuening Zhang, Fang Dai, Luojia Deng, Yijie Wu, Jianlei Gu, Qingmin Lin, Xiaolei Wang, Yan Kong, Hui Lu

## Abstract

In the realm of biomedical research, efficient data analysis and processing are crucial due to the escalating volume and complexity of data generated by research teams. Managing these vast arrays of localized data presents significant challenges, necessitating precise, efficient, and reproducible analysis methodologies to ensure the integrity and reliability of scientific outcomes. Traditional management of analysis codes, computing environments, and the inherent difficulties in result traceability due to team dynamics often lead to inefficiencies and potential risks in maintaining academic integrity. Furthermore, while online storage platforms such as Dryad, GitHub, and Docker facilitate data, code, and environment management, they do not inherently guarantee the reproducibility of results, with issues like data incompleteness, forgotten parameters, or software discrepancies posing additional challenges. To address these critical gaps, we developed a BioMedical data Analysis Platform (BMAP) to offer online and localized categorized management of research assets. BMAP enhances workflow efficiency by transforming complex pipelines into user-friendly web applications, promoting consistency and standardization across team analyses. Its comprehensive web analysis module and seamless integration with data and computing resources support automated result reproducibility and visualization. According to the assessment, 1,692 omics-related figures from 101 recent articles, across 45 visualization types, were tested with BMAP, which could cover 37.8% of the types and 64.3% of the figures. BMAP also enables the sharing and enhancement of research methods through its cloud platform, allowing researchers to utilize the previously developed and validated tools, thereby reducing redundant effort and minimizing analytical discrepancies due to methodological differences.

## Introduction

In biomedical research, the analysis and processing of data have become the core of laboratory scientific work. With the continuous growth of research team data and the rapid development of analytical technologies, researchers face substantial challenges in managing and processing vast localized data^1^. These data require not only precise and efficient processing methods but also standardized and reproducible analysis procedures to ensure the accuracy and reliability of research results^2^. However, research teams often face challenges such as managing analysis codes and computing environments, as well as issues like redundant development of functionalities due to the loss of code when team members depart, and difficulties in tracing analysis results subsequent to their departure. These issues not only consume substantial research resources but also complicate the management of academic integrity reviews.

Research teams can manage scientific process through online storage platforms, enhancing the traceability and reproducibility of results. For example, data, code, and computing environments can be stored on platforms such as Dryad(https://datadryad.org), GitHub(https://github.com), and Docker(https://www.docker.com), which play a crucial role in improving the traceability and reproducibility of outcomes. However, due to the independence of these systems, it is not possible to test if results are reproducible based on the stored data. Incomplete data storage, forgotten analysis parameters, or software version discrepancies can all lead to challenges in automatically reproducing analysis results.

To address these challenges, we have developed the Biomedical Data Analysis Platform (BMAP), which supports localized and categorized management of analysis codes, computing environments, and data. BMAP can transform complex analysis codes into visualized Web applications, reducing redundant development efforts and ensuring consistency and standardization of analyses within teams. Through its end-to-end Web analysis module and interconnectivity with data and computing environment storages, BMAP enables automated reproducibility and visualization of results. Additionally, users can publish or download other researchers’ end-to-end analysis methods on the BMAP cloud platform, further assisting and accelerating localized research.

## Results

### Complexity of biomedical data analysis

Biomedical data analysis involves intricate workflows that handle diverse data types and analytical processes. As illustrated in Figure 1, the analysis process can be divided into two main categories: omics data and medical information data.

**Figure 1.**
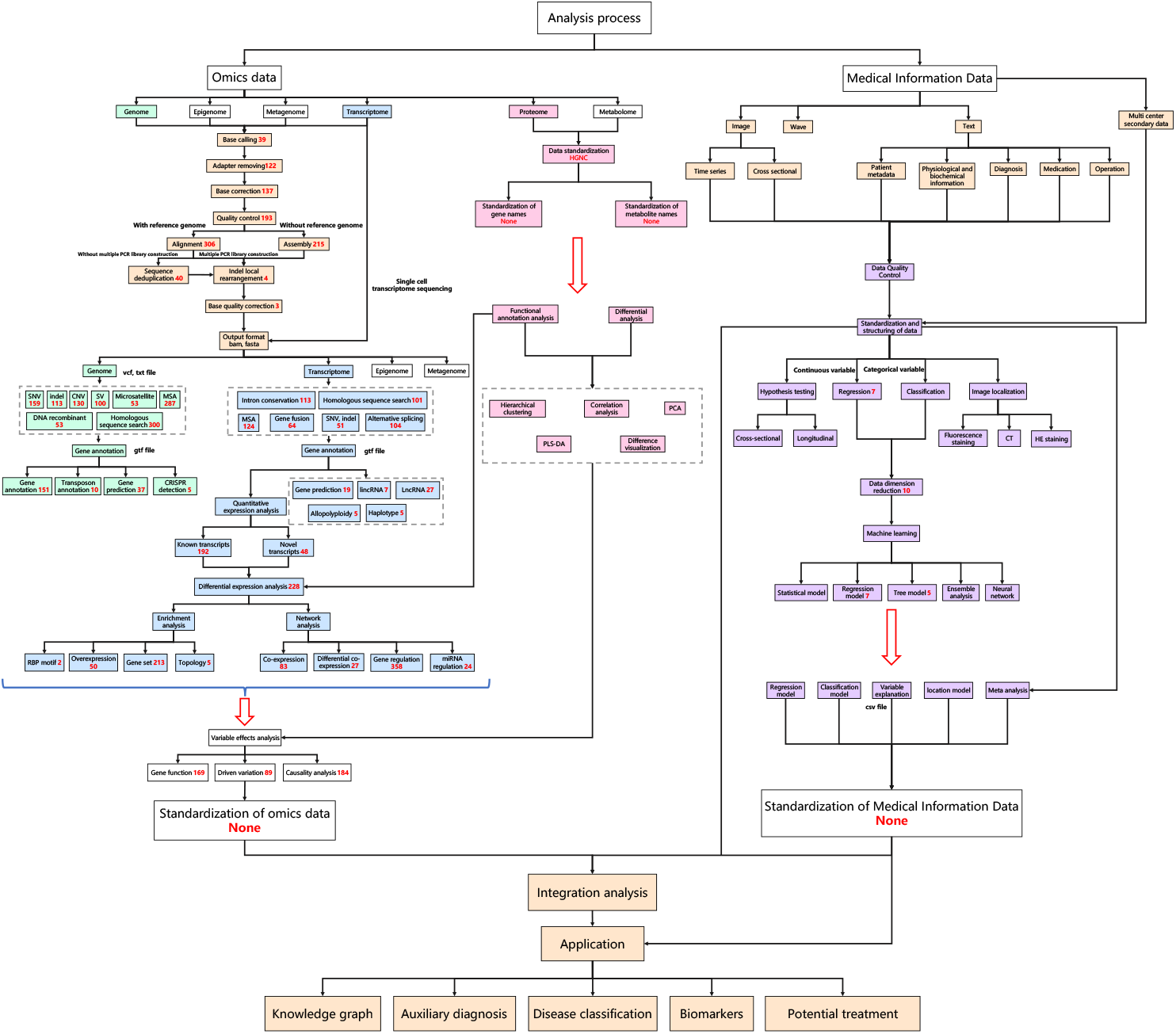
Overview of biomedical data analysis workflows. Each box in the diagram represents a specific sub-function within the biomedical data analysis workflow. The numbers inside the boxes indicate the count of available software tools that can perform the respective functions.

Each category includes multiple types of omics or different modalities of medical data, following detailed pathways to standardize and integrate the data for comprehensive analysis. At each step, the available tools also vary, ranging from a few to several hundred^3^.

The extensive array of software tools introduces dependencies and potential conflicts, making it challenging to set up a functional analysis environment. For example, in cancer genomics analysis, Figure 2a illustrates the specific workflow, including steps such as adapter trimming, sequencing quality control, sequence alignment, and variant calling. Single step can involve up to 306 different software options, resulting in a combinatorial explosion of possible software workflows exceeding 10^18^ combinations. A successful analysis case (specifying the software used for each step) involves software and computing environments that span multiple programming languages and dependencies, including 5 different file formats, 6 system libraries, 14 analysis software packages, and so on, all of which must be compatible with the underlying operating system (Figure 2b). Thus, ensuring compatibility among various tools and managing software versions are common hurdles for researchers. These complexities can significantly hinder the reproducibility and efficiency of data analyses, highlighting the need for an integrated platform to streamline these processes and reduce conflicts.

**Figure 2.**
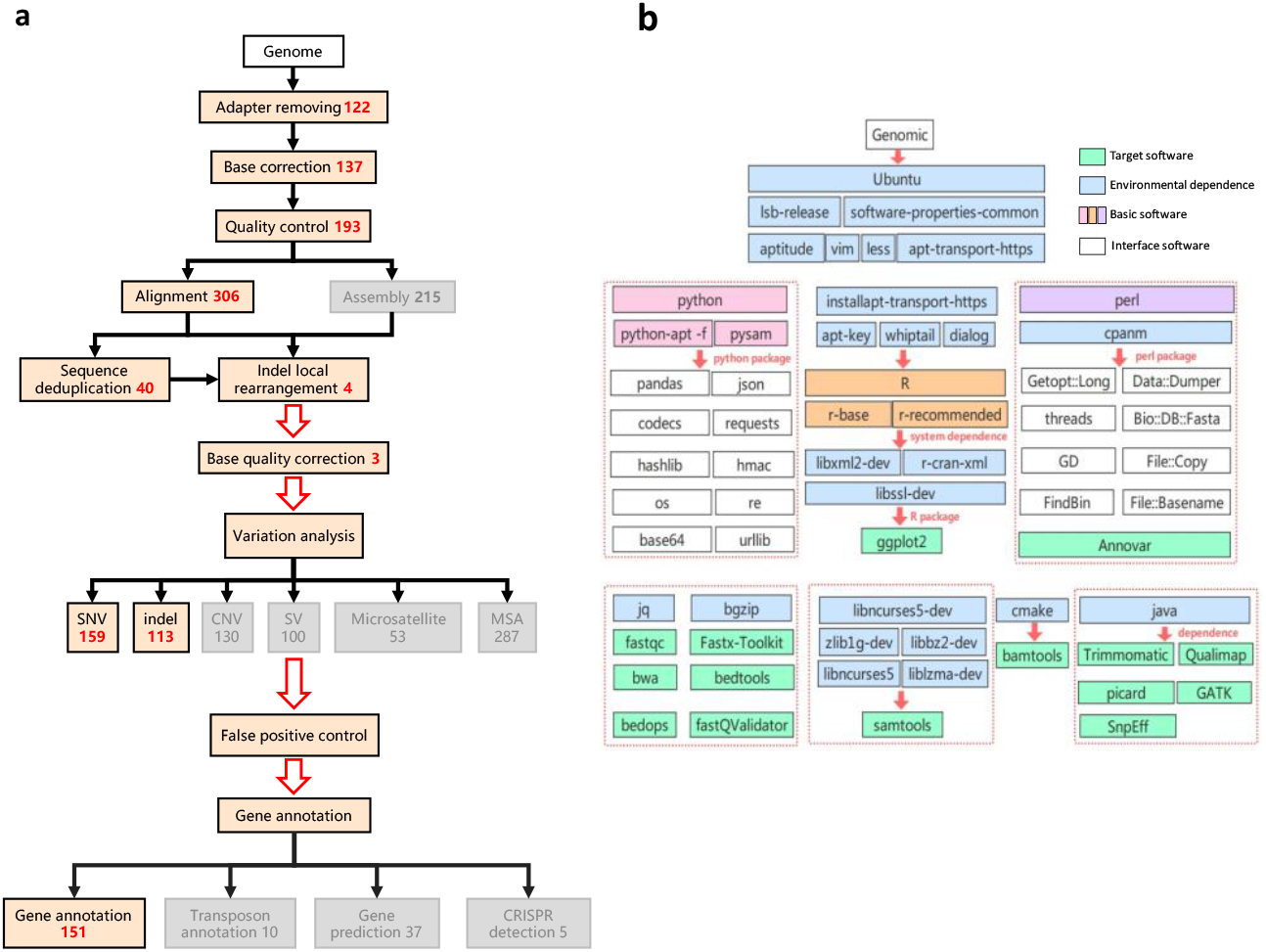
Details of cancer genomics analysis. **a**. The analytical process of cancer genomics analysis. **b**. The corresponding computational environment of cancer genomics analysis.

### Architecture

The architecture of the BMAP is illustrated in Figure 3a. BMAP is composed of a storage center, an application design system, an analysis system, a user data management center, a high-performance computing center, and an edge computing center. In the storage center, users can classify and store data, code, and computing environments (singularity images) according to their intended purpose, format, customize access permissions, and other custom categories. In the application design center, developers can specify the code and computing environment (singularity image) required to run each application, and visually edit the web operation page, including help documents, parameter configuration controls (drop-down menus, text boxes), computing resources, web result page layout, and access permissions, and then publish the designed end-to-end web application to the analysis system to be shared with other users. Researchers can install and use their own BMAP applications locally, as well as download web applications from the cloud center. Researchers can utilize their own BMAP applications locally or download and install shared applications from the publicly accessible BMAP cloud center at https://bmap.sjtu.edu.cn/.

**Figure 3.**
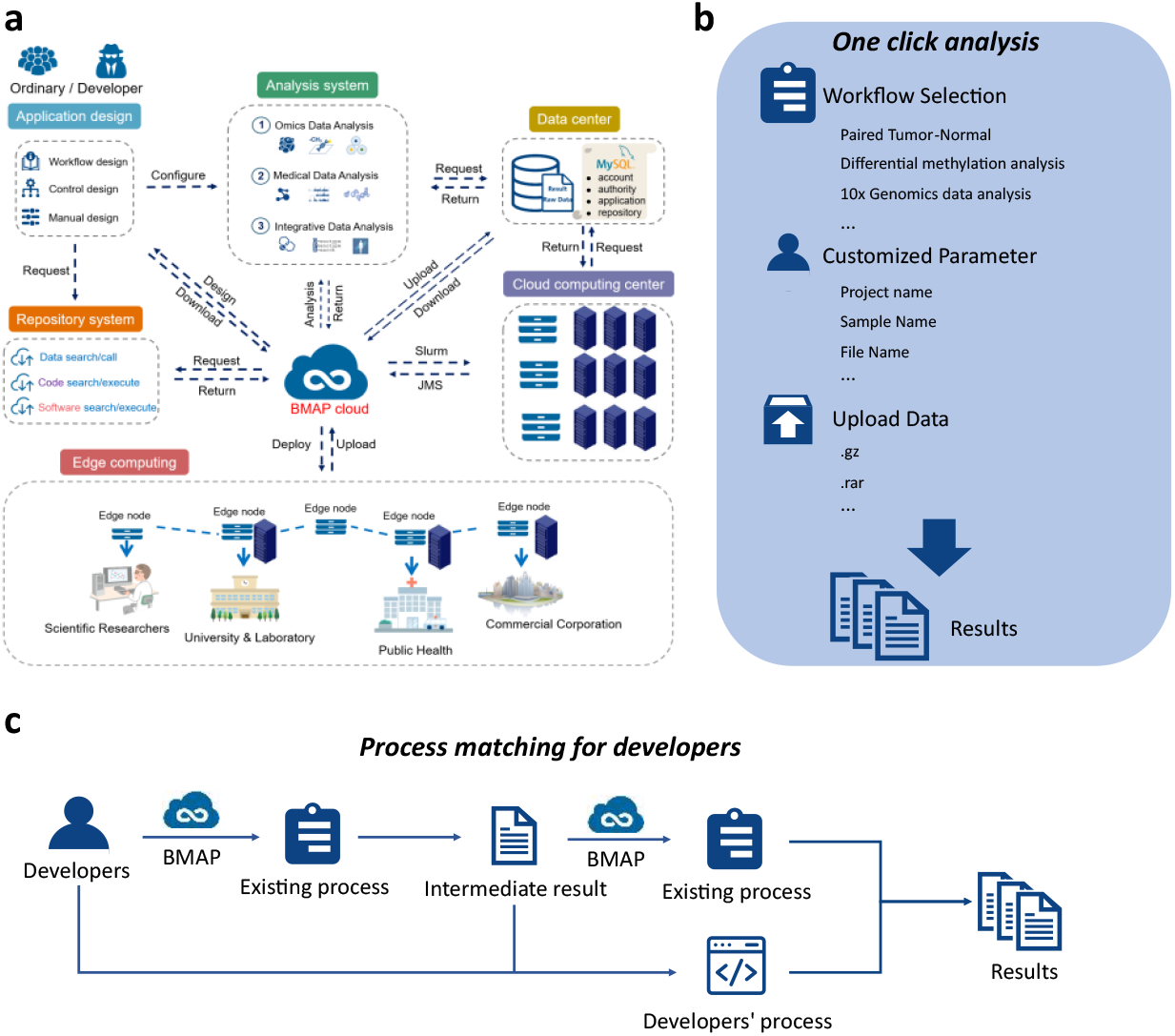
Overview of BMAP. **a**. The architecture of BMAP. **b**. The process of one-click analysis. **c**. Customizable workflow integration for developers.

For data analysis, there are two modes of use: a one-click analysis mode for all users and a customizable workflow mode for developers. As shown in Figure 3b, the one-click analysis mode allows users to select existing workflows in BMAP, set personalized parameters on the corresponding interface, upload data, and directly click to start the analysis and obtain results. This mode is user-friendly and efficient, making it accessible even to those with limited technical expertise. On the other hand, developers can leverage the customizable workflow mode to create tailored analysis workflows. They can combine any existing workflows in BMAP with their own analysis code in various forms, allowing for a high degree of flexibility and specificity in their analyses (Figure 3c). This mode supports advanced users who need to integrate bespoke computational processes, thereby enhancing the BMAP’s versatility and capability to meet diverse research needs.

### Examples of Benchmark Pipelines in the BMAP cloud center

There are currently numerous tools available for the analysis of biological data. Each tool has specific conditions under which it can be applied, resulting in a variety of pipelines with varying effects. The accuracy of the pipelines integrated in BMAP were all comprehensively evaluated. Although several research efforts have evaluated the accuracy of the latest available tools, there is currently no comprehensive evaluation of the accuracy of each pipeline integrated in the open access biomedical data analysis platform.

A standardized analysis pipeline for transcriptomic data was established in BMAP following the best guidelines in a benchmark analysis study^4^. The code and the docker image were stored in the BMAP code repository and software repository, correspondingly. To make the researchers could further evaluate the accuracy of the RNA-seq analysis pipeline, a set of RNA-seq standard data which included transcriptome sequencing data and qPCR data for approximately 959 genes across 8 samples was constructed based on the SEQC database^5^. The accuracy of RNA-seq pipeline was assessed by comparing the analysis results of the standard data with the relative gene expression values obtained by qPCR. Due to the non-normal distribution of gene expression data, the Spearman correlation coefficient was employed to measure the accuracy of the analysis process. The RNA-seq pipeline achieved a mean score of spearman score with 0.8446, and the accuracy rate was determined to be 85.3% (Figure 4 a-e).

**Figure 4.**
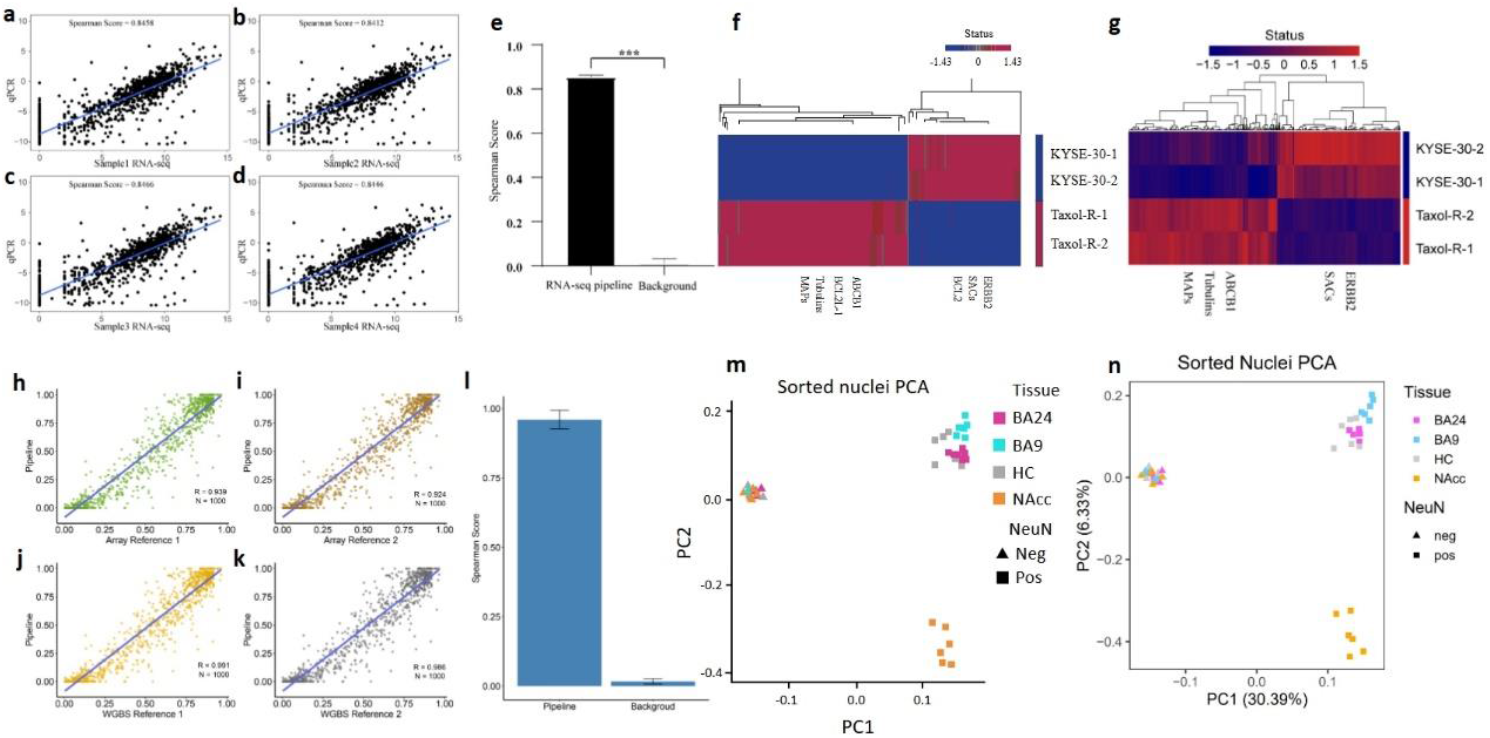
**a-d**. Spearman correlation plots comparing RNA-seq pipeline results with qPCR expression in four samples. **e**. Comparison of RNA-seq pipeline results with those obtained using randomly selected genes. **f**. The schematic diagram of Figure 1E in the original article^5^, which identified seven previously reported paclitaxel resistance-related genes. **g**. RNA differential expression analysis results automatically obtained using the method recommended by Wu, integrated within BMAP. The results show that Wu’s method, as integrated by BMAP, detected 5 out of the 7 previously reported paclitaxel resistance-related genes from panel f. **h-k**. Spearman correlation plots comparing BMAP-WGBS pipeline results with reference results. **l**. Comparison of BMAP-WGBS pipeline results with those obtained using randomly selected methylation sites. **m**. The schematic diagram of Figure 1a in the original article^8^, showing the results of principal component analysis using methylation sites detected from raw WGBS sequencing data of autosomes as input data. **n**. The method for methylation site analysis was replaced with the WGBS methylation data analysis from the BMAP Cloud Center, followed by the same principal component analysis method. The results indicate that the methylation sites detected by the BMAP WGBS analysis application did not alter the original conclusion regarding the observed clear segregation of cell types (neuronal from non-neuronal nuclei) and brain regions within the neuronal.

The pipeline was also evaluated by using a real data from previously published paper^6^. The provided heatmap illustrates the differential expression of genes in paclitaxel-resistant human esophageal squamous cell carcinoma (ESCC) cell lines (Taxol-R) as well as the transcriptome sequencing data of ESCC cell lines (KYSE-30). Compared to the original paper, there are two of seven marker genes (BCL2 and BCL2L1) were not found by using this best guidelines, the other five marker genes were detected and the relevant patterns of the differential expression were completely consistent with the original text, with an accuracy rate of 71.4% (Figure 4 f, g). This further demonstrates that due to the opacity of parameters, even using the best workflows recommended in the literature, it is challenging to fully reproduce results. However, the BMAP workflow, evaluated using standard datasets, could provide users with an objective benchmark, giving them a clearer understanding of the workflows they employ.

The standardized Whole genome bisulfite sequencing (WGBS) analysis pipeline for methylation data was established in BMAP following the guidelines in Krueger^7^ and Akalin’s^8^ studies. This pipeline utilizes input data from bisulfite sequencing (BS-Seq), and identifies CpG sites in samples through a series of steps such as quality control, filtering, alignment, and annotation. It facilitates differential analysis of methylation sites and regions between different groups, presenting users with visualized results. To assess the accuracy of the WGBS analysis pipeline, we collected a set of human prostate cancer cell line (LNCa, PrEC) datasets that were detected using both WGBS (Illumina HiSeq 2500) and microarray (Infinium MethylationEPIC BeadChip). These original datasets were downloaded from NCBI-SRA/GEO (SRP089722, GSE86833). We randomly selected 1,000 CpG sites from the over 700,000 CpG sites covered by the original dataset and performed a correlation analysis between the results of the user-customized WGBS analysis pipeline and the two reference pipelines from the original publication. The correlation coefficient indicates the accuracy of the customized software. Next, we tested the accuracy of the WGBS analysis pipeline in BMAP and obtained an average correlation of 0.960 (Figure 4 h-l). The result indicates that the released pipeline has considerable accuracy and reliability.

The accuracy of the pipeline was also evaluated by using the GSE96612 data obtained from the research conducted by Rizzardi et al^9^. The DNA methylation was examined by whole-genome bisulfite sequencing (WGBS) in neuronal and non-neuronal populations from four brain regions. The PCA plot shows the distance of the CpG methylation levels of samples in different brain regions (neuronal n = 22; non-neuronal n = 23). The original data from the paper were processed through the methylation analysis workflow at the BMAP cloud center to identify methylation sites. Subsequent principal component analysis yielded results consistent with the conclusions of the paper (Figure 4 m, n).

### Examples of Enhancing Research Accessibility and Reproducibility with BMAP

BMAP offers comprehensive support for various research projects, encompassing storage for code, software, data, and operations for visualization. For instance, all code, computing environments, and test data involved in developing the false positive variant quality control software FVC are stored in BMAP’s corresponding repositories, ensuring that research findings are always reviewable and reproducible. BMAP simplifies the process for research areas such as yeast cell counting^10^, single-cell study^11^, multi-omics study^12^, GWAS^13^, and wristwatch data analysis^14^ by providing a feature for visualization design and application publishing, which facilitates the transition from command-line to one-click visual operations. Additionally, for third-party applications like diseasGPS^15^ that have their own dedicated websites, BMAP enhances their accessibility and usability by categorizing these tools and providing navigation links.

### Analysis Coverage

For a biomedical data analysis platform, the ability to support a wide range of diverse and comprehensive analyses if crucial. Naturally, analysis coverage is a key criterion for evaluation a biomedical data analysis platform.

To assess the coverage and breadth of analyses supported by BMAP, we systematically searched PubMed for articles highly relevant to bioinformatics analysis published over a year in the journals Genome Biology, Nucleic Acids Research, and Genome Research. After screening and selection, 101 articles were included in the study. We systematically categorized and evaluated the various omics-related visualization results presented in these articles. Specifically, we assessed whether BMAP could reproduce these visualizations.

In these 101 articles, a total of 1692 omics-related figures were presented. These figures encompassed probability density plots, PCA plots, boxplots, and other commonly used bioinformatics visualizations such as heatmaps, Manhattan plots, and survival curves, covering 45 different types of visualization results. The results showed that BMAP was able to cover 37.8% of the figure types and 64.3% of the figure count. Additionally, for frequently used analyses, BMAP’s coverage was even higher (Figure 5a). For figure types that appeared in more than 40% of the articles, BMAP even achieved a coverage rate of 100%. Figure 5b specifically shows the results from articles in Genome Biology. In these 17 articles, BMAP achieved a coverage rate of 64.3% for the figure count and 50% for the figure types. These results indicate that BMAP has significant advantages and potential in supporting diverse bioinformatics data analysis and visualization, demonstrating its wide analysis coverage.

**Figure 5.**
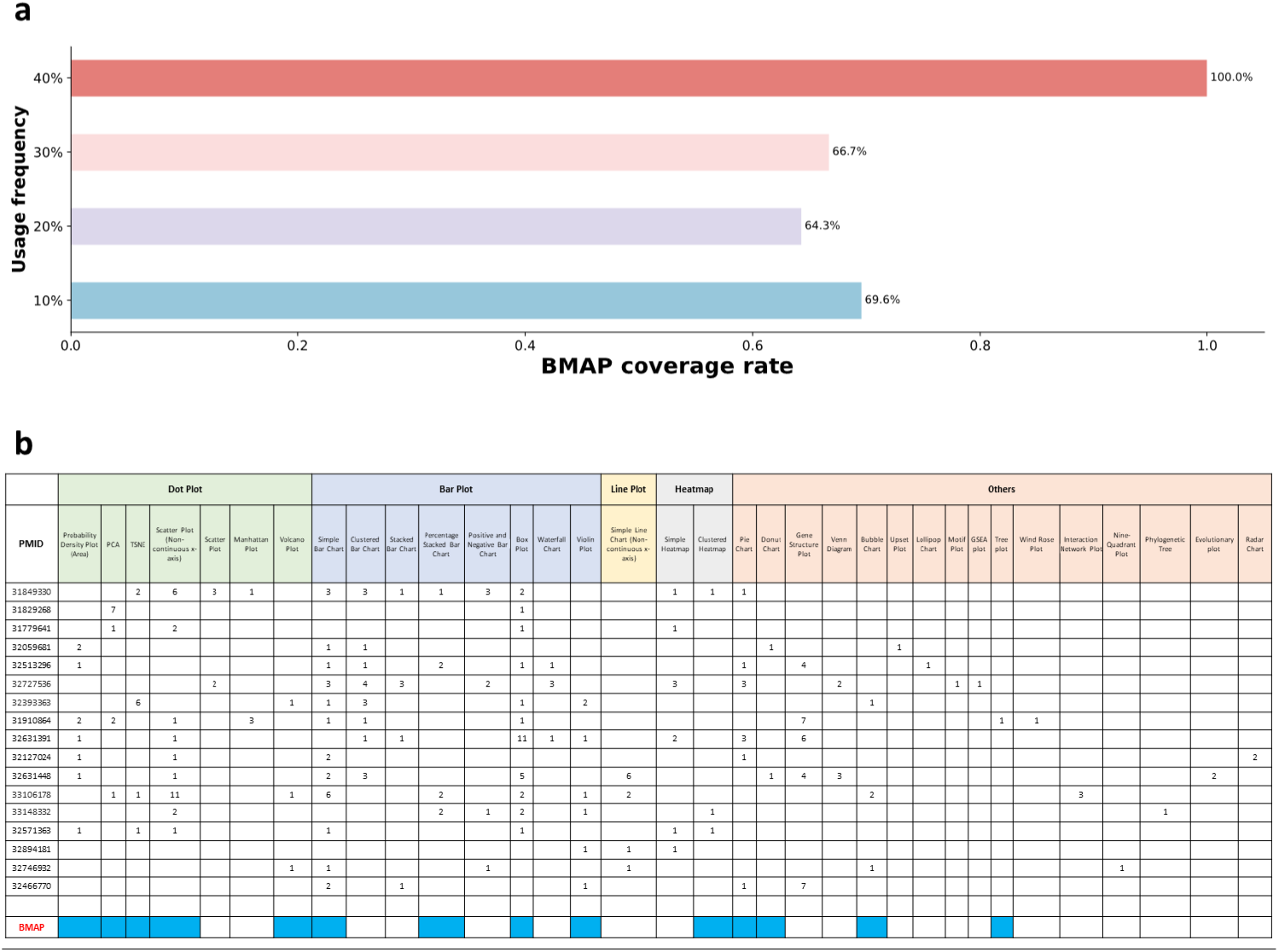
Analysis coverage of BMAP. **a**. The BMAP coverage rates across different usage frequency. **b**. Analysis coverage across articles from Genome Biology.

### Functionality Assessment and Comparison

A comprehensive biomedical data analysis platform needs to possess various capabilities to meet the diverse data processing and analytical needs in the biomedical field. We conducted a detailed assessment of the functionalities supported by BMAP and similar platforms, as shown in Figure 6. The comparison includes platforms such as TCGA^16^, SRA^17^, GitHub, Galaxy^18^, GenePattern^19^, and Clinvar^20^. The evaluation dimensions encompass 41 aspects, including data storage capabilities, analysis functions, software management, and online and localized management.

**Figure 6.**
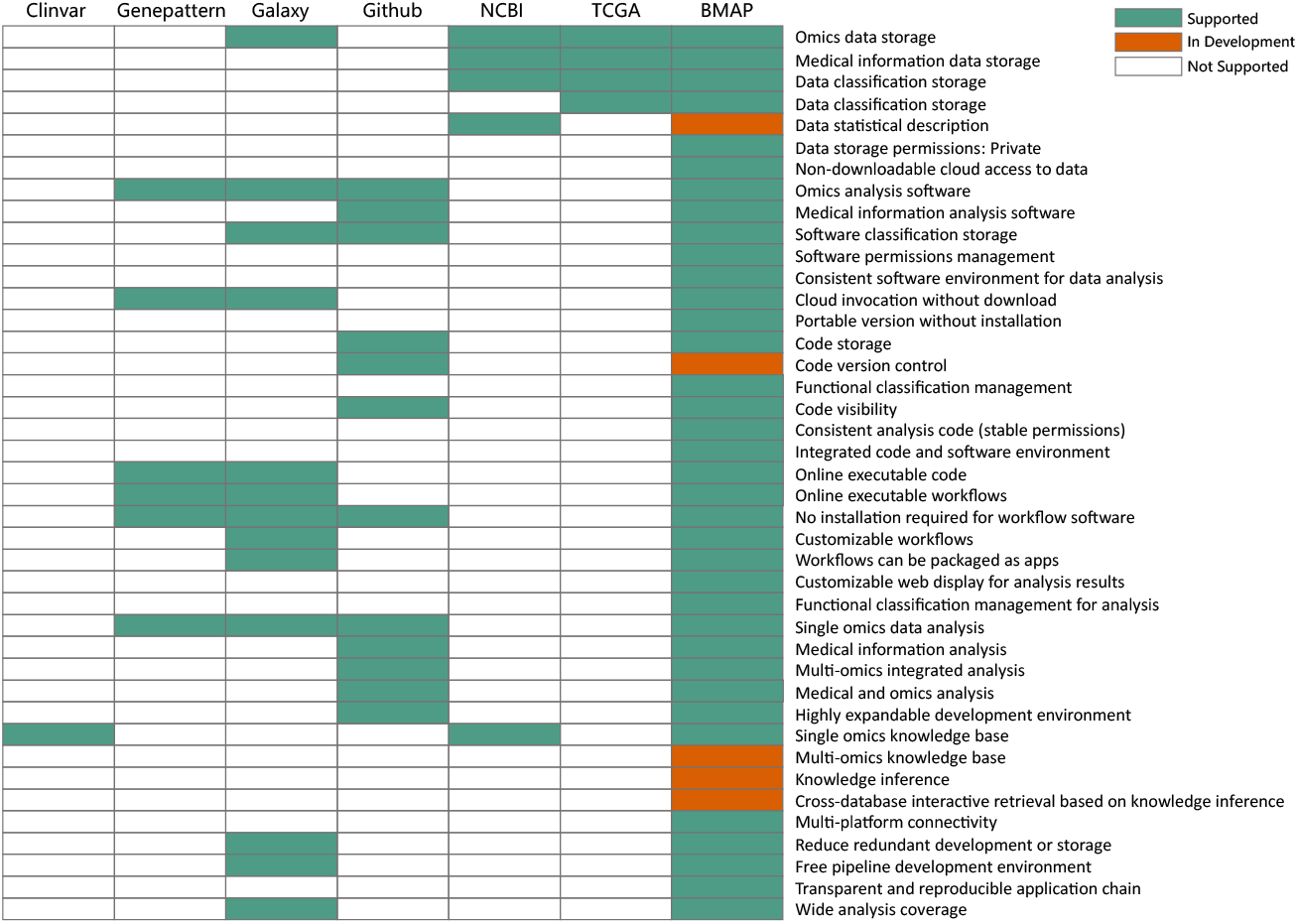
Functional comparison of BMAP with other platforms.

BMAP demonstrates a high level of support in key functionalities, including omics data storage, medical information data storage, data statistical description, and software permissions management. In contrast, other platforms offer limited support in these areas. For example, while NCBI and TCGA provide some data storage and analysis functions, they lack in software permissions management and a consistent software environment. BMAP not only provides a consistent software environment for data analysis but also supports cloud invocation without download and portable versions without installation, which is superior to GitHub. Although Galaxy and GenePattern offer online analysis tools, their functionalities in code management and version control are not as comprehensive as BMAP.

In terms of customizable workflows, BMAP allows workflows to be packaged as applications and offers customizable web displays to present analysis results. Other platforms, such as Clinvar, have weaker functionalities in customizable workflows and application packaging. BMAP also supports single omics data analysis, medical information analysis, multi-omics integrated analysis, and a highly expandable development environment. Additionally, functionalities such as knowledge inference, multi-platform connectivity, reducing redundant development or storage, free pipeline development environment, and a transparent and reproducible application chain are within the platform’s support scope. These capabilities give BMAP significant advantages in data management and analysis efficiency for research teams.

## Methods

### Reproducibility

BMAP enables users to repeat the same analysis multiple times with the same results. The application used in the analysis consists of standalone computational environment, code, and configuration parameters. The entire computing environment used in an analysis, such as dependency software and libraries, are bundled into one or more containers, which are managed by BMAP software repository and remain fixed even as software or libraries change over time. The executable code consists of the main program and configuration parameters. The main program for each application will not change over time. The configuration parameters are customized by the user, but they can be visualized in tabular form in the results, ensuring that the analysis parameters could be used and traced back for each repeat analysis.

The applications integrated into BMAP cloud must meet at least one of the three conditions to ensure the consistency with the analytical methods in published research: 1) The application is designed and released by the authors of the published article, and the code, computing environment, and parameters are same with the analytical method in the article; 2) The application has been evaluated with cross-validation gold-standard dataset, and the evaluation results can be traced back; 3) The application has been validated using the data from the published articles and evaluated for consistency with the results in the articles. If the application is granted download permissions, different users can install the same cloud-based application on the local version of BMAP, using the same code, containers, and visual interfaces as in the cloud to reduce data bias caused by differences in analysis tools.

### Accessibility

BMAP enables users to perform data analyses by providing a web-based interface for managing data and applying computational applications to analyze the data. Users can access public data or their stored private data in the BMAP data repository based on a unique identifier. Applications can non-download access the exist data according to the unique identifier in BMAP data repository or access the temporary data from the uploading web interface.

## Acknowledgements

This work was supported by the National Key R&D Program of China (2018YFC0910500); the Science and Technology Commission of Shanghai Municipality (STCSM) (Grant No. 23JS1400700); Neil Shen’s SJTU Medical Research Fund (LX, LL, HL); and the SJTU Transmed Awards Research (STAR) Grant No. 20210106 (HL). We thank Y Li, X Zhao, K Ning, B Liu, B Jin, T Ruan, X Zhang, and J Sun for helpful discussions.

